# Contextual High-throughput 3D Volume Electron Microscopy Data Acquisition Using Artificial Intelligence

**DOI:** 10.1101/2025.05.15.654255

**Authors:** Tereza Hurník Konečná, Radek Jančík, Daniela Slamková, Bronislav Přibyl, Cveta Tomova, Lolita Rotkina, Tessa Burch-Smith, Sanja Sviben, James A.J. Fitzpatrick, Kirk J. Czymmek

## Abstract

One of the greatest challenges when generating large and/or high-resolution three-dimensional (3D) volume electron microscopy (vEM) datasets is long acquisition times. We developed a method that leverages Artificial Intelligence (AI) algorithms to increase acquisition throughput for 3D datasets by creating an AI-derived mask for the targeted region of interest, referred to as Adaptive Scanning. This allowed for specific structures to be imaged at high resolution, with the surrounding area captured at lower resolution, without artificially generating or enhancing any raw data. We demonstrate that this dynamic-resolution scanning approach significantly reduced volume acquisition time across a diverse array of organisms and tissues, including brains, parasites, cultured cells, and plants. This multi-resolution strategy has the potential to enhance Focused Ion Beam Scanning Electron Microscopy and other forms of vEM, by increasing time savings by up to 2-fold or more, enabling routine generation of multiple and/or larger vEM datasets, more efficiently and cost-effectively, allowing more data collection for increase statistical power for comparative studies.

## Main

Focused Ion Beam Scanning Electron Microscopy (FIB-SEM), a form of volume electron microscopy (vEM), offers three-dimensional (3D) perspectives critical for studying complex biological structures and/or the spatial organization of organelles, cells and tissue. The Focused Ion Beam Scanning Electron Microscopy (FIB-SEM) approach is a reliable, automated process allowing for the routine acquisition of serial Scanning Electron Microscopy (SEM) images at nanometer-scale with isotropic voxels (e.g., 4-10 nm isotropic voxels) over tens of micrometers. However, the acquisition time for high-resolution 3D imaging remains a significant limitation, and imaging large volumes (on the order of tens of thousands of µm^3^) remains practically impossible on traditionally configured systems. The time required even for relatively modest volumes (e.g., 150 μm^3^ at 5nm voxel size, 600 slices @ 1.5 minutes/slice would take 15 hours) becomes a limiting factor for ultrastructural analysis of tissue volumes. Indeed, to address these limitations, vEM benefited from improved sample preparation metallization protocols ^1,2^ and/or via the application of hardware and software strategies such as “enhanced FIB-SEM”, which enabled nearly indefinite extended runs ^3,4^, serial-section electron microscopy for an entire larval zebrafish brain ^5^ or multi-beam strategies ^6^.

Artificial intelligence (AI) also has played an important and ever-increasing role in vEM studies, especially for the tedious downstream steps of annotation and segmentation for quantification and visualization of the information-rich and complex datasets ^7,8^. Likewise, in recent years, several research groups have explored solutions to optimize acquisition in SEM and enhance scanning speed for larger datasets. Indeed, various AI approaches have been employed to accelerate and refine the single-beam SEM image acquisition process and minimize sample damage. For example, the strategy of non-uniform image scanning ^9^ has been reported as an effective means to decrease scanning duration, where super-resolution reconstruction-based deep learning algorithms were applied. This strategy applies the premise of scanning low-resolution and noisy images and computationally increasing their resolution using neural networks (NN) trained on pairs of low-resolution and their high-resolution image counterparts ^10^. Alternatively, the use of random sparse scanning combined with an image based super-resolution reconstruction algorithm was applied to reduce electron dose and improve scan speed^11,12^. While effective, the previously mentioned methods are limited when the user wants to define the regions of interest (ROI) on-the-fly for high-resolution scanning while still maintaining enhanced speed or by artificially enhancing the raw data. Here, we describe a novel and robust dynamic-resolution scanning approach using AI algorithms to generate tailored masks from low-resolution images and subsequently acquiring high-resolution subsets just of the relevant pixels from the targeted ROI. Additionally, this workflow, which we termed “Adaptive Scanning” (AS), has the benefits of preserving the sample context from the low-resolution scan and greatly increasing acquisition speed, especially relevant for 3D volumes derived from 100s or 1000s of consecutive slices. Finally, AS does not generate new or enhance any acquired raw data. To achieve this straightforward workflow, we prepared our sample, established full-resolution imaging conditions, annotated the ROIs and then initialized an AI semantic segmentation model. In each subsequent image, the AI model generated a scanning mask from the low-resolution image and then applied the pixel map to only collect mask-defined high-resolution regions for the full volume. Depending on the imaging/milling parameters applied and local targeted sample features, we realized an improved time savings ranging from around 15 to over 50% compared to traditional full-resolution strategies for our test volumes. Thus, the automated AI based multi-resolution image acquisition workflow greatly improved sample throughput, especially with samples where our target structures were readily delineated and a relatively modest portion of large vEM datasets. Here, we describe the AS workflow and evaluate its robustness with diverse biological sample types and target features. We identified key parameters and robust conditions to efficiently collect more and/or larger datasets in much reduced times (and cost). Ultimately, we believe the efficiency gains with Adaptive Scanning FIB-SEM workflow (depending on research goal), reduces some of the practical barriers to collecting more, larger, higher-resolution and/or faster volume acquisitions where speed and resolution tradeoffs have limited vEM adoption and statistical power in studies.

## Results

### Adaptive Scanning Overview

The straightforward Adaptive Scanning paradigm we developed increases sample acquisition throughput by restricting the high-resolution target pixels to be collected only in AI-derived masks, which are identified from corresponding low-resolution images. In our implementation, the AS workflow is integrated in Thermo Scientific^TM^ Auto Slice & View^TM^ 5 (AS&V) Software as illustrated in the schematic overview (**Fig. 1**). First, the AS is initialized by acquiring a single low-resolution scan from the area selected for subsequent volume acquisition (**Fig. 1A**). Using digital drawing tools, the user manually annotates this low-resolution scan to identify target features and define the ROI (yellow, **Fig. 1A**). Following the initial annotation, the rest of the AS process is fully automated within the AS&V interface (**Fig. 1B**), where a low-resolution image is acquired (Low-Res, **Fig. 1B**, **Fig. 2A**) and automatically segmented by the AI algorithm into two regions: the ROI and background (AI-generated mask, **Fig. 1B**, **Fig. 2B**). The user-defined target features/objects within the ROI are selected to reflect the specific user’s needs. Depending on the scientific question under consideration, the ROI can range from organelles (e.g., mitochondria) to a whole cell within a given field of view, without limitations on the number of objects selected. The AI-generated segmentation mask of ROI is utilized for high-resolution scanning (**Fig. 2C**), resulting in a final image that merges the high-resolution targeted scan of the ROI with the low-resolution background (Merged image, **Fig. 1B**, **Fig. 2D**). Optionally, the low-resolution scans, high-resolution scans, and AI-generated masks can be saved separately for the use of efficient post-processing strategies outlined in the discussion. This procedure is repeated for each slice in the imaged volume, producing a series of images. Subsequently, the acquired data can be processed, further annotated, analyzed and/or visualized using 3D volume reconstruction tools such as Napari ^8,13^, DeepMIB ^7^, and Amira software, etc. (**Fig. 1C**, **Fig. 3**).

**Figure 1.**
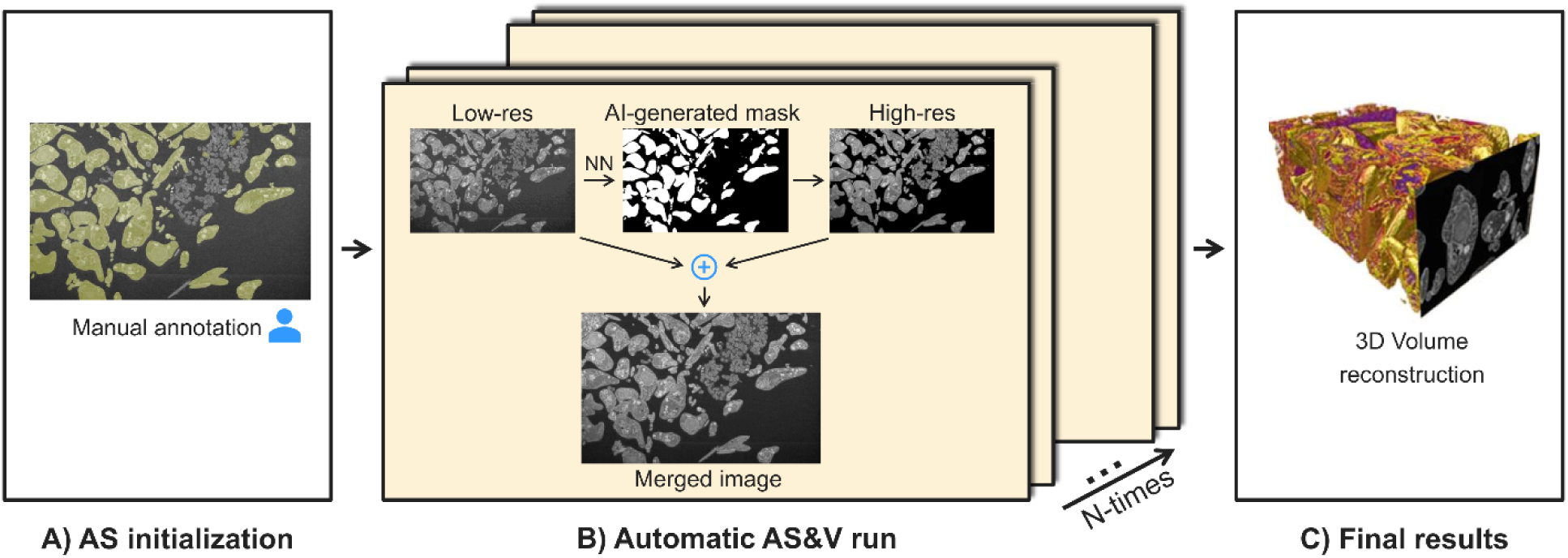
Schematic illustration of Adaptive Scanning workflow. A) During the AS initialization of the AS&V run, a single low-resolution scan is acquired from the area selected for volume acquisition, where the user manually annotates the ROI. B) The automatic AS&V run proceeds as follows: a low-resolution image is acquired, which is then automatically segmented into two regions – ROI and background (i.e., the rest of the image) – by the AI algorithm. The segmentation mask is used for high-resolution scanning. The final product is a merged image consisting of high-resolution scans of the ROI and low-resolution scans of the background. This process is repeated for each slice of the imaged volume, resulting in a series of images. C) The final results of the workflow can be optionally visualized, processed, and analyzed using a 3D volume reconstruction tool such as Thermo Scientific™ Amira Software.

**Figure 2.**
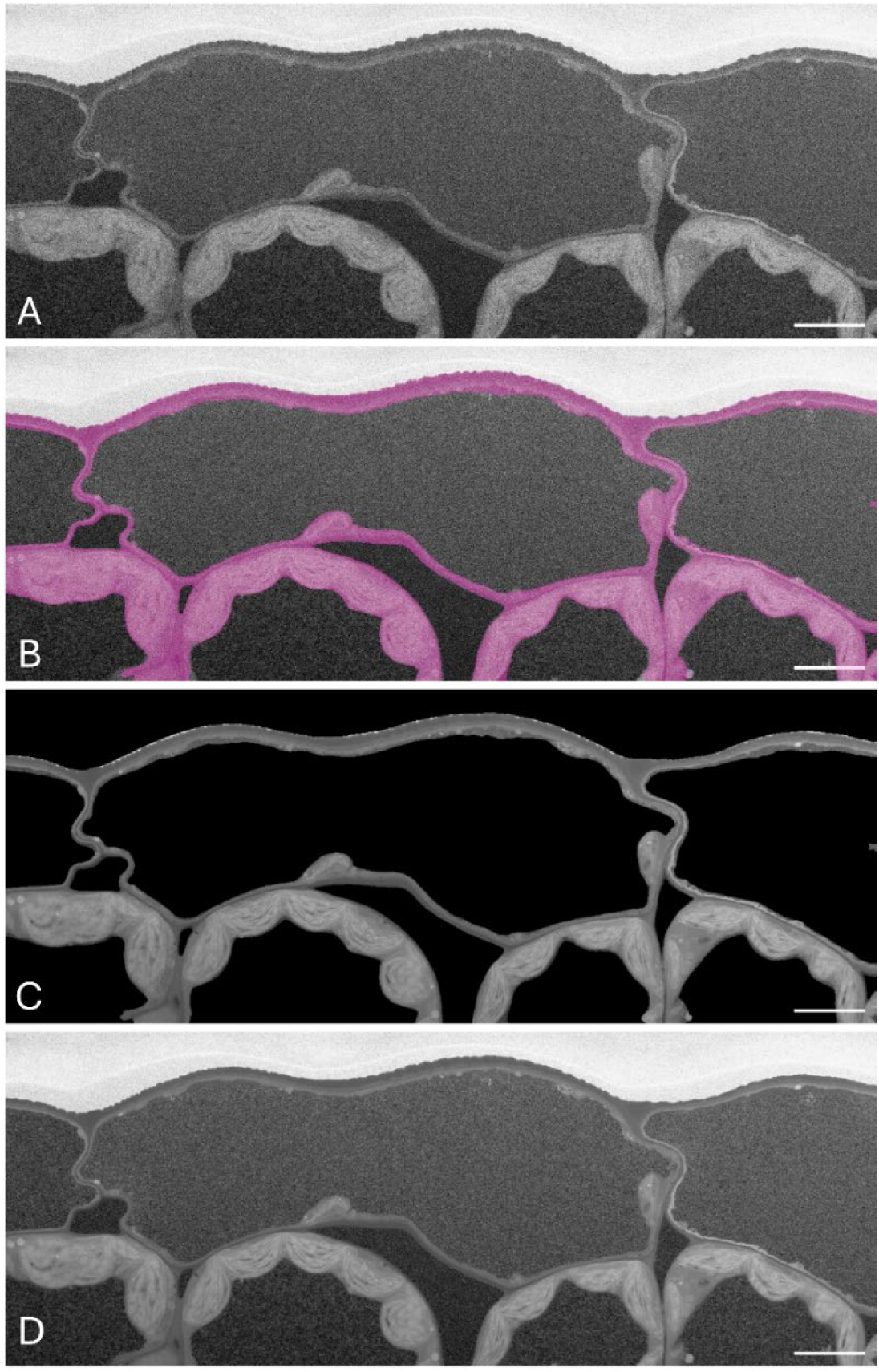
*Nicotiana benthamiana* (tobacco) epidermal cell. A) Low-resolution scan (2 keV; 0.2 nA; 1 µs; 1225×471, Voxel size 50×50×6 nm; Horizontal Field Width (HFW) 61.25 µm). B) Low-resolution image of “background” with segmented the plant cell wall and cytoplasm region for next step of scanning in high-resolution on slice 5 with segmentation mask (ROI: plant cell wall and cytoplasm region). C) High-resolution scan of ROI (2 keV; 0.2 nA; 1 µs; 10211×3928, Voxel size 6×6×6 nm; HFW 61.25 µm). D) Merged image: low-resolution “background” and high-resolution (ROI: the plant cell wall and cytoplasm region). Scale bars = 5 µm.

**Figure 3.**
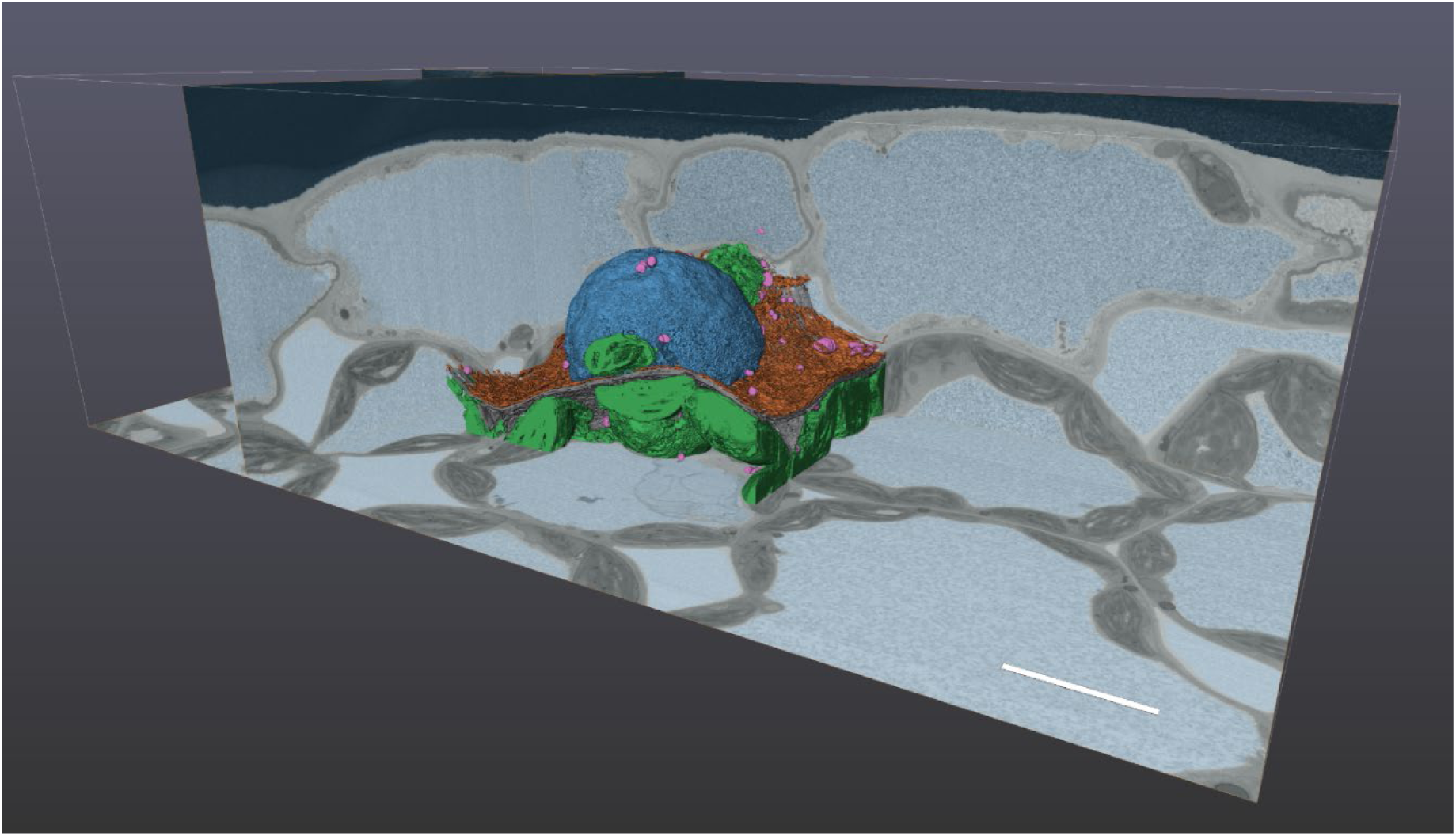
Adaptive Scanning 3D segmented sub-volume of *Nicotiana benthamiana* (tobacco) leaf epidermal cell. We demonstrate the high-resolution segmented 3D rendering of a sub-volume derived from the annotation and embedded within the full high-resolution AS volume (grey scale) and merged with the low-resolution contextual information depicting the vacuoles (light blue). The selected sub-volume showing detail of the nucleus (dark blue), endoplasmic reticulum (brown) mitochondria (pink), chloroplasts (green) segmented using AI Deep Learning in Amira software. Scale bar = 5 µm.

### Adaptive Scanning AI algorithm

The underlying AI algorithm for segmenting ROI is a semi-supervised, NN-based method that generates segmentation masks of ROIs for low-resolution images. The generated masks, which are subsequently used to target areas for high-resolution scanning, correspond to the regions identified by the AI algorithm as the most similar to those in the user’s manual annotation of the ROI.

Adaptive Scanning builds upon previous work^14^, which aimed at segmenting videos from digital cameras with an emphasis on accurate segmentation around ROI boundaries. We extended the capabilities by adapting the AI algorithm for a series of low-resolution EM micrographs. Our modifications include an adaptive mask thresholding technique, additional image preprocessing, and optimization of the NN hyper-parameters, as described in detail^14^.

The architecture of the AI algorithm is outlined in **Supplementary Fig. 1**, comprising of three main NNs: reference encoder, query encoder, and decoder. First, the information about the ROI, provided in the form of the initial scan and its corresponding manual annotation, is processed by the reference encoder and stored as deep reference features. Following this initialization, a segmentation mask for an image of a new slice is generated. The current image is processed by the query encoder into deep query features. These query features are then matched with their most similar reference features counterparts and passed to the decoder to generate the segmentation mask for the current image.

The segmentation algorithm aims to correctly classify pixels as either ROI or background. Correct classifications of ROI and background pixels are denoted as True Positives (TP) and True Negatives (TN), respectively. However, segmentation algorithms generally produce two types of errors: False Positives (FP) and False Negatives (FN). FP (i.e. over-segmentation) refers to the background pixels incorrectly classified as ROI, while FN (i.e. under-segmentation) refers to ROI pixels incorrectly classified as background.

Typically, there is a trade-off between FP and FN – improving one type of error might worsen the other. Specifically, in the AS workflow, under-segmentation leads to the loss of high-resolution image data in ROI, while over-segmentation reduces the potential gain in acquisition time. Since preserving the quality of the acquired high-resolution data in ROI is an important requirement for the reliability of AS, in this trade-off between FP and FN errors, a higher priority is set for minimizing the FN over the FP in the algorithm design. The specific trade-off was achieved by optimizing the method’s hyper-parameters using an adjustable Fβ score and implementation detailed previously^14^.

To enhance the flexibility and robustness of the AS workflow, and to address the possibility of a changing cellular landscape, it is necessary to update the segmentation algorithm during the vEM process. The AS process can be dynamically adjusted by reannotating the ROI as needed, effectively restarting the AI segmentation algorithm. This capability is particularly useful when significant changes in the sample’s structure, contrast, and/or features occur throughout the acquired volume, or when a different ROI needs to be selected. Additionally, minor segmentation adjustments, such as adjusting the ratio between over-segmentation and under-segmentation, can be easily performed within the AS&V software without the need to reannotate new ROIs or restart the AI algorithm scans.

To assess and experimentally validate the AS algorithm and workflow, and determine its capabilities and limitations in accelerating volume acquisition, we designed four experiments on data from diverse samples: 1) Determine the segmentation quality in the AI-guided ROIs, 2) Define and assess factors that accelerate volume acquisition using AS, 3) Verify the potential for whole cell large volume acquisition, and 4) Characterize the required precision of manual annotation.

### Segmentation Quality in the AI-guided ROIs

One of the critical requirements of AS is to preserve the high-quality of data within the selected ROIs. The high data quality in ROIs would be lost if the AI algorithm under-segmented the low-resolution scans, resulting in incomplete high-resolution scans of the desired ROIs. Therefore, the following experiment validated the amount of under-segmentation within ROIs during the AS process.

To evaluate the amount of under-segmentation, we employed the miss rate metric, also known as the false negative rate. The miss rate is defined as the ratio of FN pixels to the total number of pixels in the ground-truth ROI (**Eq. 1**).

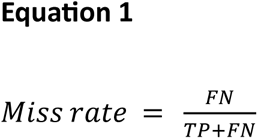

(FN = # of False Negative pixels, TP = # of True Positive pixels)

The miss rate metric can be interpreted as the proportion of the ROI area with lost image quality. According to its definition, the miss rate is independent of the relative size of ROI to the whole image. To quantitatively assess the miss rate of the AI segmentation algorithm, we acquired a dataset consisting of 8 scanned volumes of various samples and ROIs, including mouse kidney mitochondria, rat brain myelin sheath, and cultured Chinese Hamster Ovary (CHO) cells. Subsequently, we manually annotated ROIs in 31 of these images (evenly distributed along the scans) to set a ground truth for comparison with the AI segmentation masks. On these test datasets, we achieved a miss rate of 0.88%, indicating that only a minimal portion of high-quality information was lost in the ROI. This demonstrates the high reliability of AS in preserving the quality of critical data. An example comparison between the manual ground truth and the AI segmentation is visualized as single slices in **Fig. 4**.

**Figure 4.**
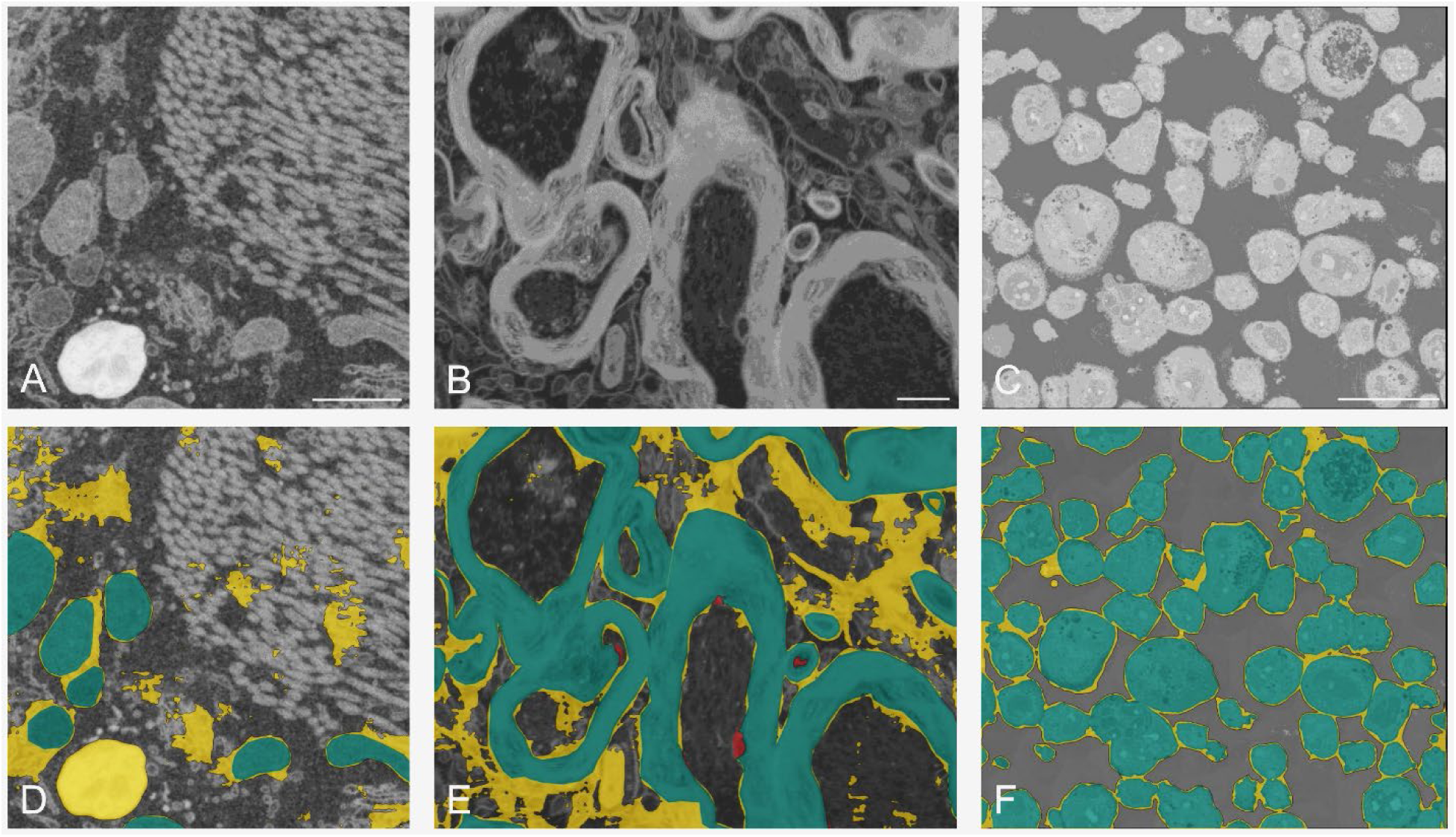
Low-resolution image and visualization of AI segmentations from different datasets: A & D) mouse kidney tissue (ROI: mitochondria); B & E) rat brain tissue (ROI: myelin); C & F) CHO cells (ROI: whole cells). TP = # of True Positive pixels, FP = # of False Positive pixels, FN = # of False Negative pixels. Aqua = TP, yellow = FP, and red = FN. Scale bar: A & B) 1 µm, C) 30 µm.

### Large Volume Acquisition

We demonstrated the capabilities of AS for extended volume acquisition by acquiring a large dataset of 6114 slices from a single cell of a *Nicotiana benthamiana* (tobacco) leaf epidermal cell, where each cell has a very large central vacuole and numerous airs spaces that are off-target but constitute a considerable portion of each image (**Fig. 2**, **Fig. 3**, EMPIAR-11831). The initial setup, including the manual annotation of ROI on the first slice, is shown in **Fig. 5 A-C** and **Suppl. Fig. 2**. In this experiment, to ensure ROIs are segmented optimally, the AS was manually readjusted four times on slices: 508, 1198, 2702, and 2805. Following the final adjustment, over 3000 consecutive slices were acquired fully automatically without any user intervention, and the segmentation masks remained consistent with the target ROI (**Suppl. Fig. 2**). **Fig. 5D, G & J** display selected low-resolution scans from this experiment, along with their corresponding segmentation masks (**Fig. 5E, H & K**) and high-resolution scans (**Fig. 5F, I & L**). This illustrated the robustness and adaptability of the AS workflow in maintaining volume acquisition over an extended series of slices.

**Figure 5.**
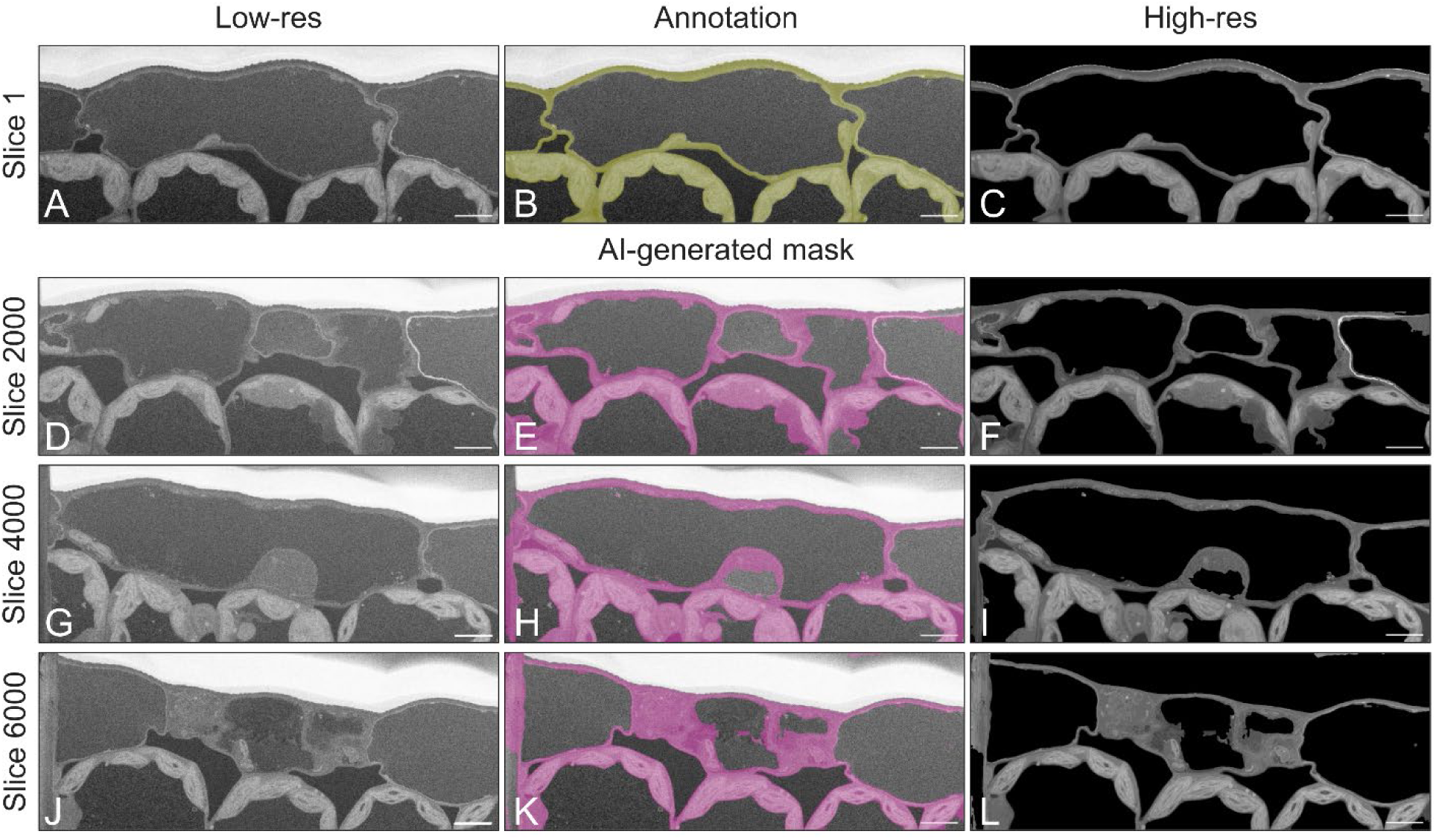
Example of a long AS run with re-annotation. *Nicotiana benthamiana* (tobacco) leaf epidermal cell. A, D, G, & J) Low-resolution scan of slice 1, 2000, 4000, and 6000 respectively (2 keV; 0.2 nA; 1 µs; 1225×471, Voxel size 50×50×6 nm; Horizontal Field Width (HFW) 61.25 µm). B) Annotation of ROI (the plant cell wall and cytoplasm region) in the first low-res image for next step of scanning in high-resolution on slice 1. C) High-resolution scan of ROI (2 keV; 0.2 nA; 1 µs; 10211×3928, Voxel size 6×6×6 nm; HFW 61.25 µm). E, H, & K) Low-resolution scan of slice with the AI-generated mask for slice 2000, 4000, and 6000 respectively. F, I, & L) High-resolution scan of ROI of slice 2000, 4000, and 6000 respectively (2 keV; 0.2 nA; 1 µs; 10211×3928, Voxel size 6×6×6 nm; HFW 61.25 µm). Scale bars 5 µm.

### Accelerating Volume Acquisition using Adaptive Scanning

To assess the increased throughput using AS, we analyzed the tobacco leaf dataset described in the previous experiment. The average time-per-slice was calculated from a sub-series of 200 slices using both AS-based and conventional acquisition methods, allowing us to calculate the time saving of AS as follows:

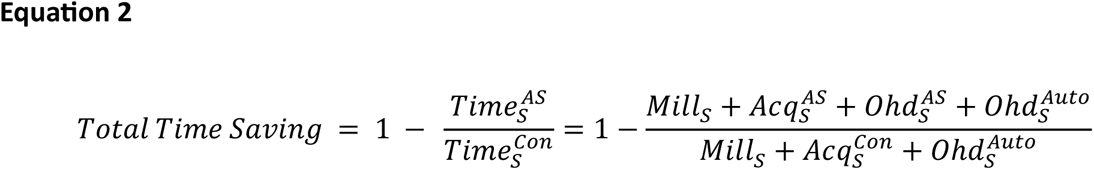

**Definitions:** S (Slice) AS (Adaptive Scanning), Con (Conventional Scanning), Mill (Milling), Acq (Acquisition), Ohd (Overhead), Auto (Automation)

The total time saving achieved for the whole cell dataset was 49%, while the acquisition time saving (excluding the milling time) was 57%. For a dataset of 6144 slices, conventional acquisition would have required approximately 17 days and 17 hours of microscope time, whereas AS reduced this to less than 8 days and 22 hours.

To understand the time-saving potential of AS, it is crucial to review the relationship between key parameters in a typical FIB-SEM experiment. The entire volume acquisition time depends on the milling time, image acquisition time, and overhead components, which include all indirect steps intrinsic to the data collection process (e.g., image drift correction, frequency of auto-functions, stage movement, etc.). The image acquisition time saving when using AS depends on the ROI percentage within the imaged area, the actual high- and low-resolution values, as well as the dwell time. The dependencies between each component comprising the data collection process can then be compared (**Fig. 6**). The milling time directly impacts the total time required for volume acquisition and thus also the total time saving (**Fig. 6**). To demonstrate this, we calculated four scenarios where the image size and acquisition parameters were identical, but the milling time per slice varied from 15 seconds to 2 minutes. This resulted in a linear decrease in the total time saving, from 53% to 36%. Thus, milling time is an important parameter influencing the total time saving for a given volume. It should be mentioned that different milling settings will affect the overall time per volume regardless of whether conventional scanning mode or AS mode was applied. However, combined with AS, milling time becomes a more powerful factor contributing to the total time saving for identical volumes.

**Figure 6.**
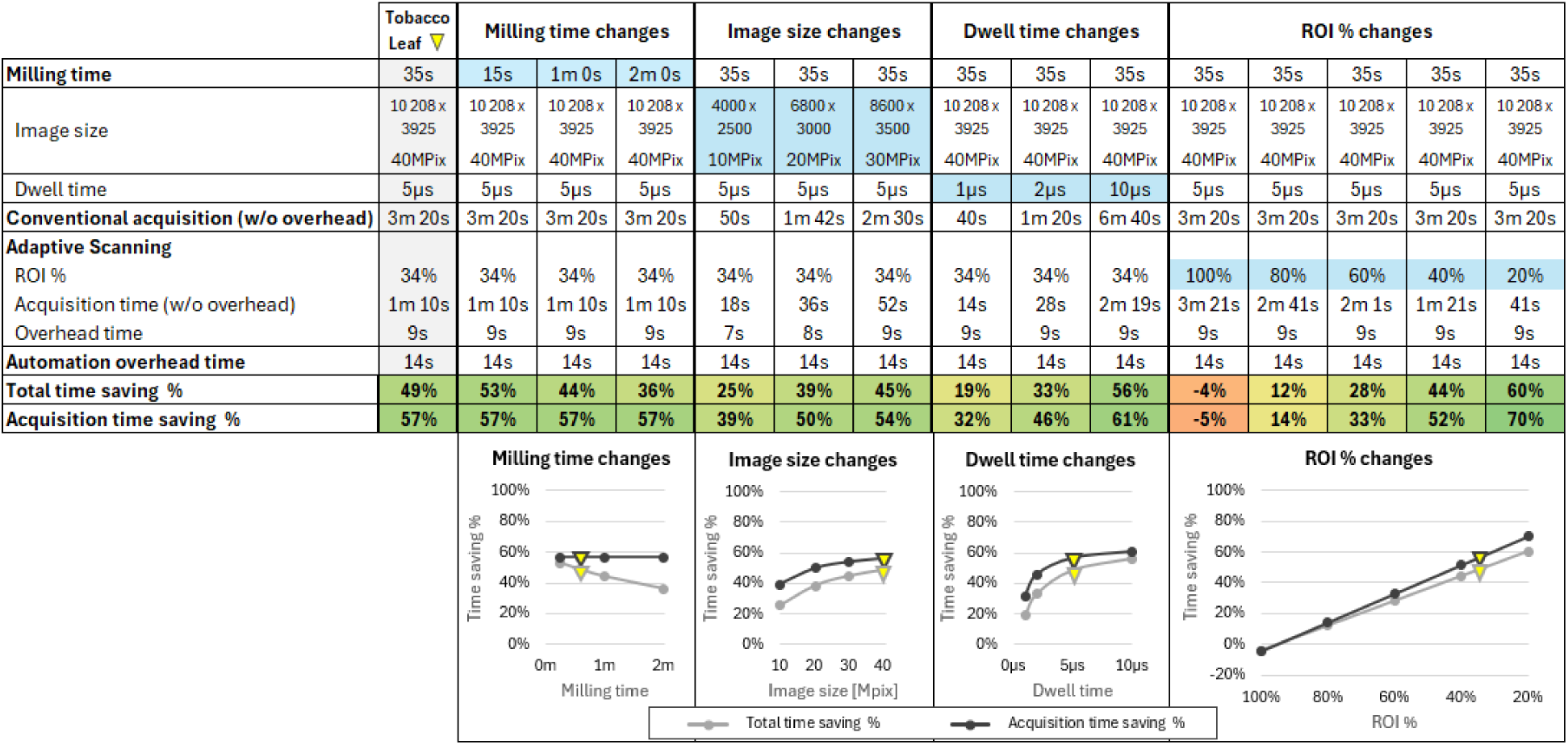
Factors Influencing AS Time Savings. This Table and graphs show how the time saved is affected by several factors comprising the data collection process. The first column (grey) labeled as Tobacco Leaf (Fig. 3**, and Suppl. Table 1 & 2**) represented our experimental acquisition where the same sample was acquired under conventional (full-resolution) and AS modes. The acquisition time saved was calculated according to **Eq. 1** based on the log file of the automation software. To illustrate the influence that each parameter had on the time saved we then altered only one parameter (Milling Time, Image Size, Dwell Time, ROI Area – light blue cells respectively) and kept all other settings identical.

The percentage of ROI within an image can significantly contribute to image acquisition time saving. The examples ranged from 20% to 100% of interesting pixels acquired at identical scanning settings (**Fig. 6**). These examples represent realistic scenarios where ROIs can be as low as 20%, e.g. individual cells withing a resin block, or up to 80% where much of the entire field-of-view is of scientific importance. As expected, the dependencies between ROI percentage and saved time are linear, with lower ROI percentages within the imaged volume resulting in higher time-saving effects. Another critical parameter influencing the time saving when using AS is the image size. The AS approach is especially advantageous when higher image size values are required for the ROI. Although each parameter contributes differently to the time saving, it is crucial to consider the cumulative effect of all parameters. Therefore, when evaluating or optimizing the time saving using AS, it is important to assess all relevant parameters comprehensively.

### Precision of Manual Annotation

Another aspect that might influence the AS process — both the quality of data and the increased acquisition efficiency — is the manual annotation of the ROI during the setup. In this experiment, we tested three different annotations on the same dataset and compared the resulting AI segmentation masks (**Suppl. Fig. 3**). We examined the influence of annotation precision on the AI-generated mask accuracy using the whole-cell tobacco test dataset. On the low-resolution scan of the first slice (**Suppl. Fig. 3A**), we created three different manual annotations: optimal (**Suppl. Fig. 3B**), over-annotation (**Suppl. Fig. 3C**), and under-annotation (**Suppl. Fig. 3D**). The optimal annotation contained an ROI (cell wall and cytoplasm region) bounded by clear contrastive edges. The over-annotation was a thicker annotation of the cell wall and cytoplasm region, extending into the interior and exterior of the cells. The under-annotation contained the cell wall, only a few pixels wide. For each of these annotations, the AI segmentation algorithm generated masks for the acquired scans of 6114 consecutive slices.

The AI-generated masks on scans of two selected slices (**Suppl. Fig. 3E & I**) are compared in **Suppl. Fig. 3F-H & J-L**. Visual inspection of the AI segmentation for the over-annotation (**Figs. 3G, K**) indicated that the AI-generated masks spread into the background regions over time. This phenomenon could be attributed to the over-annotation not aligning well with the AI algorithm, which was trained to segment ROIs with well-defined boundaries. Without contrastive edges at the ROI boundaries, the AI-generated masks diverged from the original annotation, searching for contrastive edges in adjacent areas. Conversely, the under-annotation might be too thin, lacking sufficient visual information for the AI segmentation algorithm to accurately segment the ROIs, leading to under-segmented regions (**Suppl. Fig. 3 H, L**). The optimal annotation was confirmed to be ideal for the AI algorithm, being sufficiently wide and containing contrastive edges at the boundaries on the annotated ROI (**Suppl. Fig. 3 F & J**). Thus, the AI algorithm was able to segment this ROI well throughout the whole dataset (**Fig. 5 E, H & K**).

In conclusion, this experiment demonstrated the importance of precise initial annotation of the ROI and confirmed that the AI algorithm performs best in segmenting features bounded by edges of high contrast.

## Discussion

The primary goal of our work was to develop the integrated AI-driven application Adaptive Scanning to significantly enhance the potential of vEM imaging workflows by improving acquisition efficiency while maintaining image quality and sample context. Although the current implementation was limited to FIB-SEM, other vEM platforms such as serial block-face SEM and array tomography as well as large 2D maps could be adapted and benefit in a similar fashion. It is worth noting that generating a precise segmentation model requires both an investment of time in annotating a small dataset and expert knowledge of the target. The AS approach described here is more flexible because it does not require off-line pre-training, and the ROI annotation is performed only once at the beginning of volume acquisition. Additionally, it is agnostic to the type and shape of the ROI. Other future implementations of AS could improve the precision of the AI segmentation algorithm by allowing users to annotate more than just one image. In this setting, a user could collect tens of images of the specific sample, annotate the specific ROI, and the AI would be trained on such data, automatically segmenting the specified ROI. Further, we envision a multi-resolution AS, which would allow us to track multiple ROIs using different acquisition settings or in a more sophisticated implementation varying pixel resolution with masks to scan with different quality, pixel dwell-time or by applying interpolation.

While it increased dataset size by including low-resolution, mask, high-resolution and merged low/high-resolution images, we found the multiscale aspects of AS to be very powerful. As an example, our annotated plant cell strictly targeted the plant cell wall and cytoplasm for the high-resolution ROI, which reflected the biology we sought to measure. To maintain the integrity of the data, the AS method was designed as a comprehensive approach where the entire imaged area is scanned, combining both low and high-resolution information. This is especially important when the segmentation is not perfectly precise due to sample feature/staining characteristics that can lead to potential loss of information. Additionally, while the tissue context from low-resolution scans provided valuable information about adjacent target features (e.g. and leaf air spaces and vacuoles) these areas often have low or no cell features (and a corresponding lack of conductive staining), thus tended to be more prone to charging and beam damage artifacts. The low-resolution scans can help reduce or mitigate these issues for more stable runs and improved post-processing and segmentation. While we largely worked with the multiscale merged low-resolution/high-resolution dataset (**Figs. 3 & 5**), processing the high-resolution images only has an advantage as they are already essentially pre-segmented and thus much more constrained as far as complexity of features for further visualization annotation with AI based training. While not implemented here, it is also possible to incorporate a compression scheme applied to the high-resolution scan datasets that would greatly reduce the entire volume by compressing the number of non-interesting pixels where there is an absence of data. Accordantly, this would reduce the required disk space, memory, GPU and computational requirements of the full resolution dataset, which potentially will be a challenge with much larger whole cell vEM datasets anticipated with the more efficient AS-based workflow.

AS can be particularly beneficial for improving the ability to generate sufficient quantitative data. Details such as number, size, location, or distribution of ROIs in each volume can be the focus for generating improved opportunities for statistical validation and comparative studies of cell populations and/or experimental conditions. By creating a library of different vEM datasets where the high-resolution ROI can be reviewed within their tissue context and quickly compared across multiple 3D volumes for accurate validation of treatment components or understanding the intricate changes of physiological stress or pathology in tissues. Thus, the ability afforded by the AS strategy to capture more and/or larger high-resolution volumes demonstrated herein has the potential to allow the possibility of whole cell and quantitative statistical analysis with vEM to be more practical. Finally, generating such precise AI segmentation models of multiple ROIs would be especially beneficial in scenarios where identical samples are subject to research. In such cases, these models can be re-used multiple times, significantly enhancing efficiency. Further, considering FAIR practices ^15–17^, it is conceivable that pre-trained segmentation models be made available and accessed through ROI segmentation libraries to different research groups as shared resources, reducing the need for redundant model training.

## Methods

### Sample Preparation

The methods mentioned above were validated on the following samples: rat brain and mouse kidney tissue, Chinese hamster ovarian cells (CHO), Trypanosoma cells, and *Nicotiana benthamiana* leaf tissue. Sample preparation for FIB-SEM and AS&V with Adaptive Scanning parameters are summarized in **Supplementary Tables 1 & 2**, respectively.

Rat brain and mouse kidney tissue samples were fixed as described previously ^18^. Briefly, tissues were fixed in 2.5% glutaraldehyde, 2% paraformaldehyde with 2 mM calcium chloride in 0.15 M sodium cacodylate buffer and then post-fixed with a modified osmium tetroxide-thiocarbohydrazide-osmium tetroxide (OTO) protocol^19^, with 2% OsO4 for 1.5 hour, 2.5% potassium ferrocyanide for 1.5 hours, 1% thiocarbohydrazide for 2 hours and 2% OsO4 for 4 hours. Fixation steps were followed by 1% uranyl acetate for 12 hours at 4 ◦C and lead aspartate at 60°C for 4 hours, dehydrated in a graded series of acetone and embedded in HardPlus Resin 812. The stained and embedded samples were trimmed, mounted on an aluminum stub and coated with a thin gold layer prior to FIB-SEM imaging. CHO cell suspensions were pelleted at 500G for 1 minute and washed in phosphate-buffered saline (PBS), fixed for 1 hour with 2.5 glutaraldehyde in 0.1M PBS at 37◦C, pelleted and resuspended in 2% agar and post-fixed with 2% osmium tetroxide in 0.1M PBS on ice in the dark for 1 hour. Samples were then washed in PBS, followed by water and stained with aqueous 1% uranyl acetate for 30 minutes, washed in double distilled water, dehydrated in a graded series of ethanol and transitioned to propylene oxide (PO) and Durcupan resin for infiltration and embedment (3:1, 2:1, 1:1 and 100% PO:Durcupan) and polymerized 48 hours at 60°C.

A *Trypanosoma cruzi* cell culture was spun down, and the pellet resuspended in a fixative containing 2.5% glutaraldehyde and 2% paraformaldehyde prepared in a cell culture medium. Post-fixation cells were washed 3 times for 10 minutes each in cold 0.15 M cacodylate buffer containing 2 mM calcium chloride, pH 7.4 and then incubated in a solution of 2% osmium tetroxide and 1.5% potassium ferrocyanide in cacodylate buffer for 1 hour in the dark. After incubation, cells were washed 5 times for 3 minutes each in ultrapure water and then incubated for 20 minutes in a 1% thiocarbohydrazide solution, rinsed 5 times for 3 minutes each in ultrapure water, and thereafter incubated in 2% osmium tetroxide for 30 minutes in the dark. The samples were then again washed 5 times for 3 minutes each in ultrapure water and incubated in 1% uranyl acetate at 4°C overnight. The following day, the samples were washed 5 times for 3 minutes each in ultrapure water and incubated in 20 mM lead aspartate solution for 30 minutes at 60°C. Post-incubation, the samples were washed 3 times for 10 minutes each in ultrapure water, dehydrated in a graded anhydrous acetone series (30%, 50%, 70%, 90%, 100% 3x) for 10 minutes in each step and infiltrated with Durcupan ACM resin (Sigma).

Tobacco samples were prepared as described previously^20^ based on a modified protocol^1^. Briefly, tobacco leaves (∼5 mm x 5 mm) were fixed in 2% paraformaldehyde, 2% glutaraldehyde, 0.01% Tween-20, 0.05% Malachite green in 0.1M sodium cacodylate buffer (pH 7.4) at 4^◦^C overnight. Samples were then rinsed, fixed in 1% osmium tetroxide 0.1M sodium cacodylate buffer for 4 hours followed by treatment with 2.5% potassium hexacyanoferrate(II) trihydrate in 0.1M sodium cacodylate buffer, 1% thiocarbohydrazide and subsequently 2% osmium tetroxide. Following fixation, samples were sequentially *en bloc* stained with aqueous uranyl acetate and lead aspartate before dehydrating in graded series of acetone and propylene oxide before embedment in a hard formulation Quetol 651 (Electron Microscopy Sciences).

### FIB-SEM Imaging and Adaptive Scanning

All samples were mounted on aluminum pins, rendered conductive via silver epoxy and/or metal coating and prepared and imaged on a Helios 5 Hydra CX DualBeam Plasma FIB-SEM (Thermofisher Scientific) or Helios 5 Hydra CX DualBeam FIB-SEM with focused plasma beams for site preparation (Xenon) and automated serial section milling (Oxygen) followed by the imaging, using the in-lens detector (TLD) secondary and/or backscattered electrons. The milling and imaging conditions for each dataset provided herein are detailed in **Supplementary Table 2**.

For Adaptive Scanning, which is integrated into AutoSlice&View workflow (Thermo Scientific^TM^ Auto Slice & View^TM^ 5 (AS&V) Software) was first collected at low-resolution image (e.g., 50×50 nm pixel size) for user manual annotation the regions of interest to train the AI segmentation network and generate the scanning mask for high-resolution imaging. Next, the AI semantic segmentation network adjusts the mask based on the subsequently acquired low-resolution image(s) and applies the image adapted mask to collect data only at the appropriate pixels for each slice to generate a high-resolution 3D image stack (**Fig. 1**). The training annotation was refined until reliable capture of the target regions and was occasionally adjusted within the sample volume if local changes in sample features/contrast warranted.

### Segmentation & Rendering

For Visualization and Deep learning segmentation in Amira 2024.2 (Thermo Fisher Scientific), a sub-volume of high-resolution data cropped out and used for segmentation. The volume loaded into the “Segmentation+” workroom and three ROIs on three different slices were selected. All voxels within each ROI were manually labeled into one of six labels: Cell exterior, chloroplast, ER, lipid, nucleus and cell wall. The AI assisted module within Segmentation+ workroom was then used to train a VGG16 UNet using all the default settings. Small islands (noise) were removed with the “Remove Islands” tool and some minor corrections were made by hand to achieve the final segmentation.

## Data availability

vEM datasets and segmentation models have been deposited in EMPIAR: *Nicotiana benthamiana* epidermal cell acquired conventional by AS&V (EMPIAR-12662; https://doi.org/10.6019/EMPIAR-12662), *Nicotiana benthamiana* epidermal cell acquired by AS (EMPIAR-12659; https://doi.org/10.6019/EMPIAR-12659), Trypanosoma cruzi cell (EMPIAR-12661; https://doi.org/10.6019/EMPIAR-12661), Chinese Hamster Ovary cell line (EMPIAR-12660; https://doi.org/10.6019/EMPIAR-12660), Rat brain tissue (EMPIAR-12675; https://doi.org/10.6019/EMPIAR-12675), and Mouse kidney tissue (EMPIAR-12674; https://doi.org/10.6019/EMPIAR-12674).

## Code availability

Adaptive Scanning is detailed in the following U.S. Patents: U.S. Patent for Data acquisition in charged particle microscopy Patent (Patent # 12,223,752 issued February 11, 2025) and U.S. Patent for Adaptive specimen image acquisition Patent (Patent # 11,982,634 issued May 14, 2024).

## Acknowledgements

Rat Brain and Mouse Kidney sample courtesy of Marie Vancová, Laboratory of Electron Microscopy Biology Centre of the ASCR České Budějovice. We acknowledge the BC CAS core facility LEM supported by MEYS CR (LM2023050 Czech-BioImaging). CHO sample courtesy of Core Facility Cryo-electron Microscopy and Tomography of CEITEC Masaryk University Brno. We acknowledge imaging support from the Advanced Bioimaging Laboratory (RRID:SCR_018951) at the Danforth Plant Science Center and use of the Thermo Scientific Helios 5 Hydra DualBeam PFIB-SEM acquired through generous donor support to the Donald Danforth Plant Science Center. We also thank Ondřej Machek and František Jeřábek, Thermo Fisher Scientific, for their contribution to Adaptive Scanning. We would like to specially acknowledge and thank Jessica Heebner and Remi Blank Thermo Fisher Scientific for Amira processing and segmentation and preparation of **Fig. 3**.

---

**Supplementary Figure 1.**
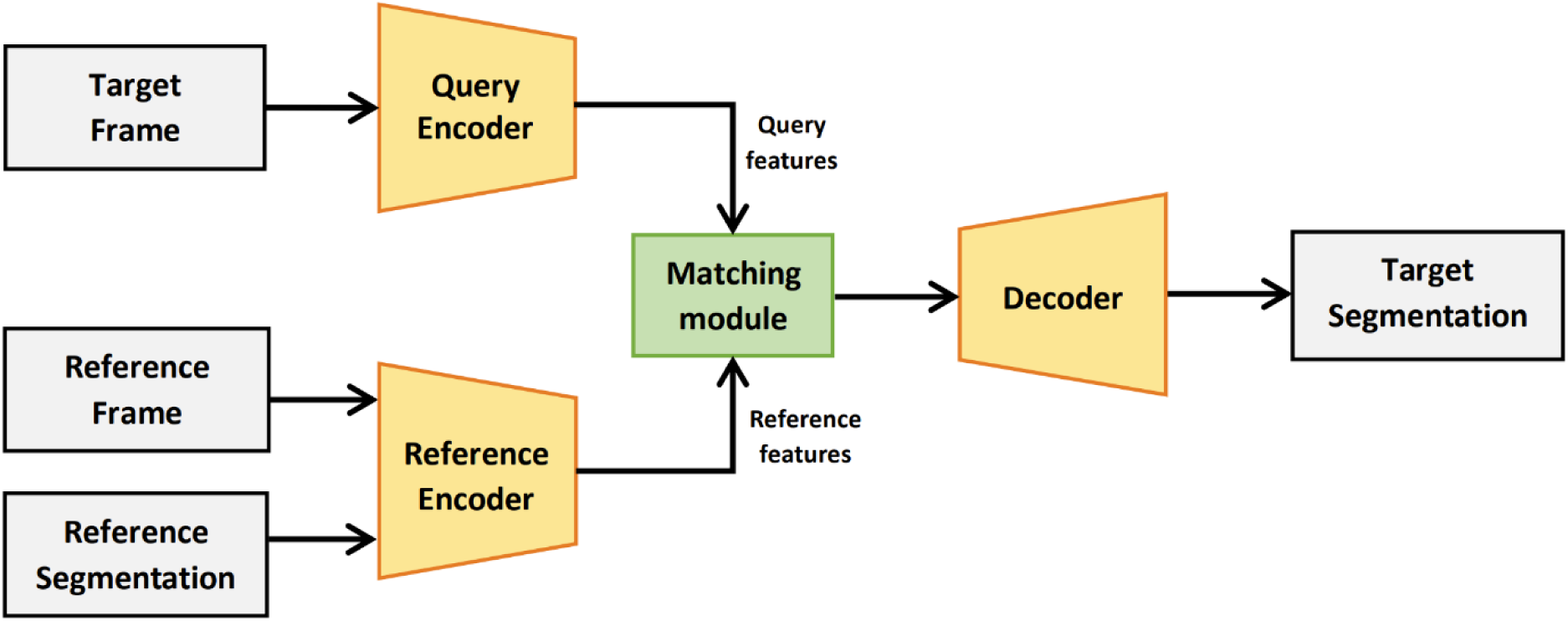
Overview of the AI segmentation algorithm architecture. It consists of several NNs (Query Encoder, Reference Encoder, and Decoder). During the AS initialization, the Reference Frame and its corresponding Reference Annotation are processed by the Reference Encoder into deep Reference features, encoding the visual information for the user-defined ROI. After slicing, a low-resolution image (Target Frame) is processed by a Query Encoder into deep Query features. These Query features are combined, in the Matching module, with the ROI information from the Reference features. The result of the Matching module is passed into Decoder, producing the final Target Segmentation. Image used with permission from Radek Jancik (Jancik 2023).

**Supplementary Figure 2.**
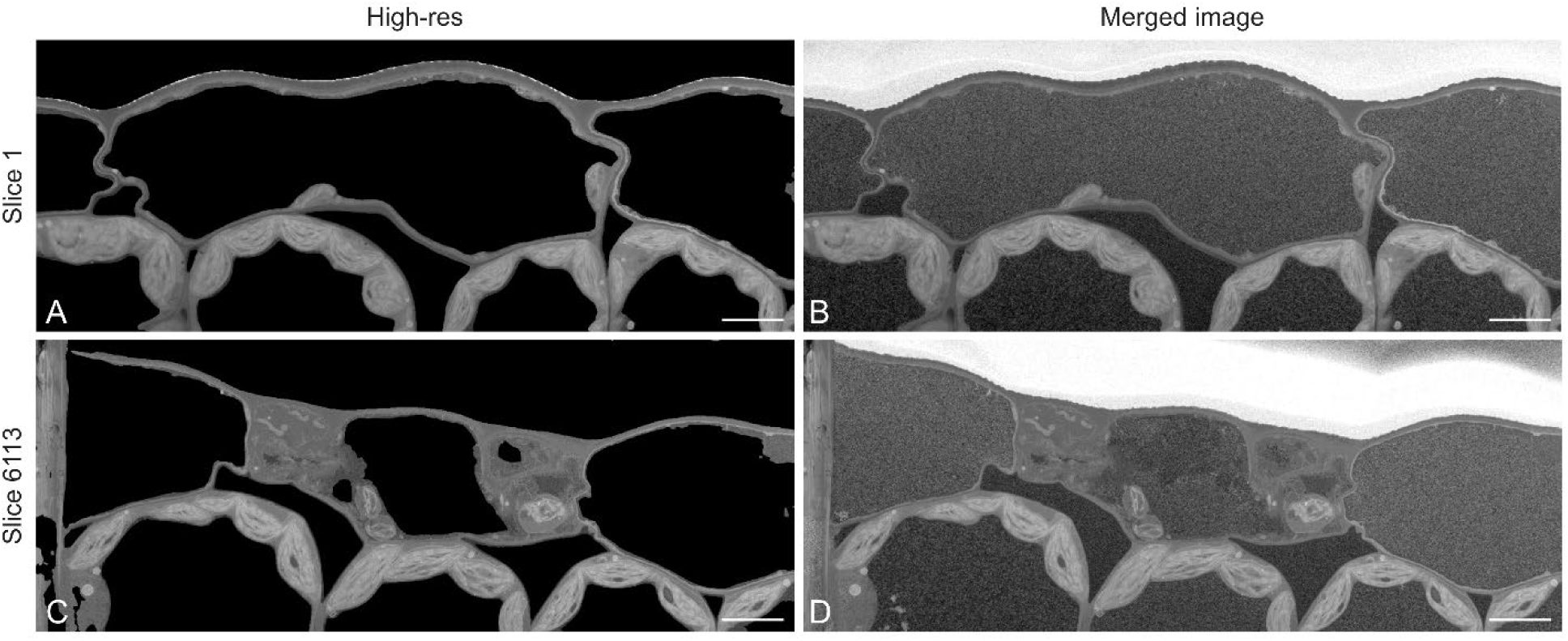
*Nicotiana benthamiana* (tobacco) leaf epidermal cell. Comparison of the high-resolution (A & C) and merged images (B & D) of adaptive scanning: low-resolution “background” and high-resolution (ROI: the plant cell wall and cytoplasm region) on 1^st^ slice and last slice (6113^nd^) of the AS&V job (2 keV; 0.2 nA; 1 µs; 10211×3928, Voxel size 6×6×6 nm; HFW 61.25 µm). Scale bars = 5 µm.

**Supplementary Figure 3.**
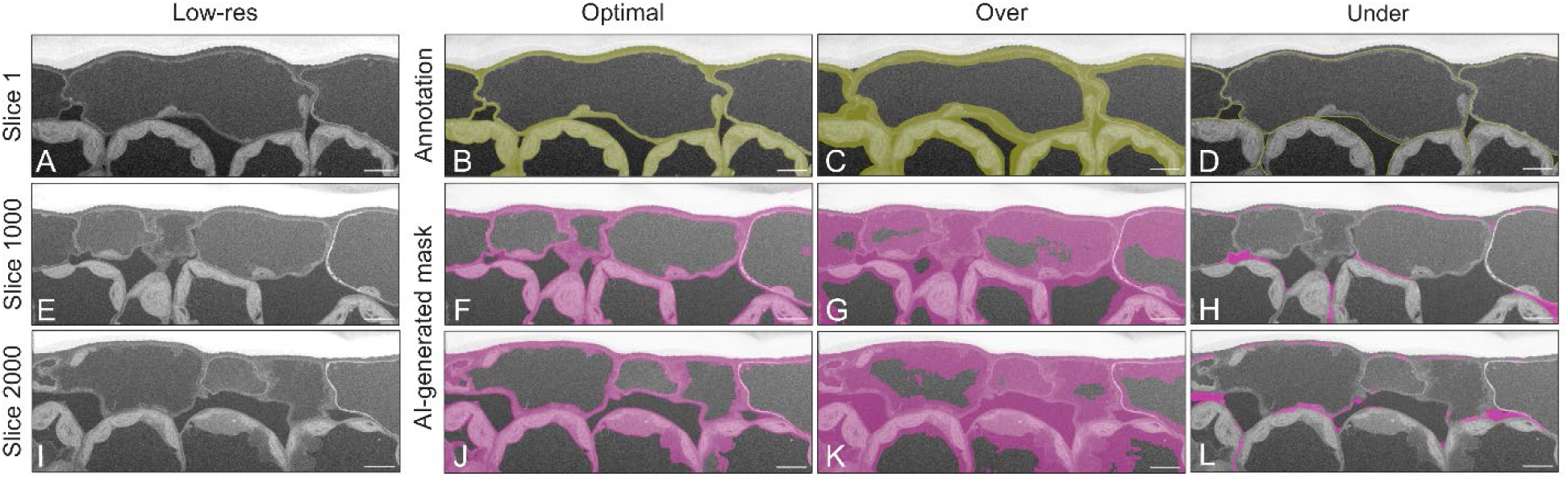
The influence of user annotation on the AI-generated output. *Nicotiana benthamiana* (tobacco) leaf epidermal cell. A, E, & I) Low-resolution images of slice 1, 1000, and 2000 respectively (2 keV; 0.2 nA; 1 µs; 1225×471, Voxel size 50×50×6 nm; Horizontal Field Width (HFW) 61.25 µm). B) An example of an optimal manual annotation. It is wide enough and is bounded by contrastive edges (ROI: the plant cell wall and cytoplasm region). F & J) The AI-generated mask for slice 1000 and 2000 respectively using the initial annotation from image B). C) An example of an over-annotation; too many features, which are not desired to be tracked, are annotated. This annotation will likely result in tracking too many irrelevant features, slowing the acquisition process. G & K) The AI-generated mask for slice 1000 and 2000 respectively using the initial annotation from image C). D) An example of an under-annotation; too narrow features were selected to be tracked. Such features might be too difficult to be tracked, likely leading to unstable segmentation, resulting in losing relevant data. ROI: the plant cell walls. H & L) The AI-generated mask for slice 1000 and 2000 respectively using the initial annotation from image D). Scale bars = 5 µm.

