## Supplementary Table 2 for "Contextual High-throughput 3D Volume Electron Microscopy Data Acquisition Using Artificial Intelligence"

### Supplementary Table 2. ASV and Adaptive Scanning Image Acquisition Parameters

| Figure # | Data collection Type | Plasma Type for Serial Milling | Milling Plasma Beam Voltage | Milling Plasma Beam Current | ASV Workflow | Application file for milling | Z-Resolution (slice thickness), (nm) | Electron Beam Voltage (kV) | Electron Beam Current (pA) | Electron Beam Dwell Time (us) | Detector Type/Mode | Detector Settings | Low/Regular Scan pixel size (X nm x Y nm) | Regular Scan Line integration | Regular Scan Frame integration | Adaptive Scanning pixel size (X nm x Y nm) | Line integration | Frame integration | ASV Estimated Image Time | Image Dimension (um x um) | Total number of slices |
|---|---|---|---|---|---|---|---|---|---|---|---|---|---|---|---|---|---|---|---|---|---|
| *Figure 1* | AdSc | Galium | 30 kV | 1.2 nA | cross-section | Si-ccs | 5 | 2 | 200 | 10 | TLD, BSE | N/A | 25 x 25 | 1 | 1 | 10 x 10 | 1 | 1 | N/A | 3807 x 2402 pixel | 488 |
| *Figure 2, 3, 5, 6 Supplementary Figure 2 &3* | AdSc | Oxygen | 30 kV | 1.7 nA | cross-section | Si, Si Multipass | 6 | 2 | 200 | 3 | TLD, Custom | 10.0 V, Mirror -15.0 V | 50 x 50 | 1 | 1 | 6 x 6 | 1 | 1 | 0:01:50 | 9375 x 3925 pixels or 56.250 um x 23.550 um | 6114 |
| Figure 4A | SEM | Oxygen | 30 kV | 0.61 nA | cross-section | Si-ccs | 3 | 2 | 200 | 5 | ICD | N/A | 3 x 3 | 1 | 1 | N/A | N/A | N/A | N/A | 4346 x 3948 pixel | 377 |
| *Figure 4B* | SEM | Oxygen | 30 kV | 0.23 nA | cross-section | Si-ccs | 5 | 2 | 200 | 5 | TLD, BSE | N/A | 5 x 5 | 1 | 1 | N/A | N/A | N/A | N/A | 2048 x 1768 pixel | 720 |
| *Figure 4C* | SEM | Oxygen | 12 kV | 64 nA | spin mill | Si - rectangular | 70 | 2 | 200 | 2 | CBS | N/A | 90 x 90 | 1 | 1 | N/A | N/A | N/A | N/A | 1536 x 1324 pixel | 183 |
| Figure 6 | SEM | Oxygen | 30 kV | 1.7 nA | cross-section | Si, Si Multipass | 6 | 2 | 200 | 5 | TLD, Custom | Suction - 10.0 V, Mirror -15.0 V | 6 x 6 | N/A | N/A | N/A | 1 | 1 | 0:03:23 | 10208 x 3925 pixels or 61.248 um x 23.550 um | 200 |
