## Supplementary Table 1 for "Contextual High-throughput 3D Volume Electron Microscopy Data Acquisition Using Artificial Intelligence"

**Supplementary Table 1. Sample preparation parameters for FIB-SEM samples**

| Figure # | Sample Type | Sample Preparation Method/ Protocol | Reference/Publication | Milling/Imaging Microscope | Software version/Mode | Sputtering for Large Area | Ion/Plasma Milling Source for the Area Preparation | Ion/Plasma Milling Source for the Actual ASV | GIS Type for Sample Area Protecting Layer |
|---|---|---|---|---|---|---|---|---|---|
| *Figure 1* | *Trypanosoma cruzi* | OTO protocol and *en bloc* staining | | Thermofisher Scientific Helios 5 UX | Thermo Scientific Auto Slice & View (ASV), version 5.10 with AI-Driven Adaptive Scanning | Iridium coating - 6 nm | Galium 30 kV | Galium 30 kV | Pt GIS |
| *Figure 2, 3, 5, 6, Supplementary Figure 2&3* | *Tobacco leaf (Nicotiana benthamiana)* | OTO protocol and *en bloc* staining | *Wickramanayake and Czymmek, 2023* | Thermofisher Scientific Helios 5 Hydra CX DualBeam Plasma FIB-SEM | Thermo Scientific Auto Slice & View (ASV), version 5.10 with AI-Driven Adaptive Scanning | Carbon coating - 10 nm, Irridium deposition - 6 nm | Xenon, 12 kV , Xenon 30 kV, Oxygen 30 kV | Oxygen 30 kV | Pt GIS, T = 45 C° , "Pt dep" |
| *Figure 4A* | *Mouse kidney* | OTO protocol and *en bloc* staining | Sheikh et al. 2023 | Thermofisher Scientific Hydra Bio UX | Thermo Scientific Auto Slice & View (ASV), version 5 | Gold coating - 20 nm | Xenon 30 kV | Oxygen 30 kV | W Multichem |
| *Figure 4B* | *Rat brain* | OTO protocol and *en bloc* staining | *Sheikh et al. 2023* | Thermofisher Scientific Hydra Bio UX | Thermo Scientific Auto Slice & View (ASV), version 5 | Gold coating - 20 nm | Xenon 30 kV | Oxygen 30 kV | W Multichem |
| *Figure 4C* | *Chinese Hamster Ovary cell line* | Osmium protocol and *en bloc* staining | | Thermofisher Scientific Hydra Bio UX | Thermo Scientific Auto Slice & View (ASV), version 5 | Gold coating - 20 nm | Xenon 30 kV | Oxygen 12 kV | None |
